# Transcriptome differentiation in *Cryptomeria japonica* trees with different origins growing in the north and south of Japan

**DOI:** 10.1101/2025.02.22.639673

**Authors:** Tokuko Ujino-Ihara, Kentaro Uchiyama, Seiichi Kanetani, Yoshihisa Suyama, Yoshihiko Tsumura

## Abstract

*Cryptomeria japonica* is a coniferous species widely distributed throughout Japan and therefore adapted to various environments. To seek genes involved in the local adaptation of this species, individuals with different origins growing at three common gardens located in the south, central and north of Japan were subjected to transcriptome analysis. The transcriptome assembly, guided by whole-genome sequence of *C. japonica*, resulted in 77,212 transcripts from 56,203 genes. Individuals were clustered into three genetic groups based on single nucleotide polymorphisms (SNPs) detected in 12,389 genes among them. Weighted gene co-expression network analysis (WGCNA) identified 25 gene modules. Comparison of representative gene expression patterns for each co-expression gene module with genetic differentiation predicted by SNPs revealed that one module exhibited a negative correlation and another a positive correlation across all three common gardens. While defense response genes were highly expressed in individuals from the Pacific Ocean side of Japan (omote-sugi), terpenoid metabolism genes were more expressed in individuals originating from the Sea of Japan side (ura-sugi). These results suggest that local adaptation associated with the alteration of gene regulation occurred in biotic stress response genes in *C. japonica*.

## Introduction

Being sessile and long-lived organisms, trees adapt to fluctuating environments through changes in their genomic DNA. Due to genetic differentiation, the response of individuals to their surrounding environment is not uniform, even within the same species. Understanding such intraspecific genetic diversity has become important for determining the vulnerability of tree species to predicted future climate change; therefore, extensive research has been conducted to identify genetic differentiation that evolved for the adaptation to local environments. Common garden trials, in which individuals originating from different natural distribution areas are planted at the same test site, are useful for researching this topic.

Previous studies have employed common garden experiments to investigate the phenotypic responses of tree lines, and, more recently, their associations with genetic differentiation (i.e., sequence polymorphisms of genomic DNA) have also been surveyed [1–3]. These studies revealed that even wind-pollinated species such as conifers, which are characterised by low or continuous genetic differentiation, showed clear phenotypic variation along geographic or climatic clines. Furthermore, next-generation sequencing technology has enabled field transcriptome analysis of tree species grown in common gardens. Differences in the transcriptome often lead to phenotypic differences [4]. Although RNA expression can be highly variable under field circumstances, recent advances in analytical tools can associate the variation of gene expression with the phenotypic traits of source populations or climate characteristics of their region of origin [5,6]. Traits related to environmental adaptation often involve the expression of multiple genes. Weighted correlation network analysis (WGCNA) is widely used to detect such co-expressed gene networks and their association with the phenotype [7]. WGCNA has been successfully applied to plant species to find key gene candidates involved in the modulation of phenotypes [8–10].

*Cryptomeria japonica* is a coniferous species distributed in Japan and southeastern China. In Japan, the species are broadly distributed from the Aomori Prefecture (40.7333°N) to Yakushima Island in the Kagoshima Prefecture (30.2500°N) [11]. Phenotypic divergences between the populations established on the Pacific Ocean side and those on the Sea of Japan side have been reported in needle morphology [12], terpene concentration and constituents [13,14], clonal propagation [15], and growth patterns [16]. These findings suggested the presence of two main *C. japonica* groups: omote-sugi (distributed on the Pacific Ocean side) and ura-sugi (distributed on the Sea of Japan side). DNA polymorphisms also supported this genetic differentiation and suggested the existence of two additional genetically diverged groups, one originating from the northern peripheral region and the other from Yakushima Island [17,18]. The phenotypic and genetic differentiation observed in the above-mentioned studies indicated that *C. japonica* is locally adapted, and thus the genes involved in this adaptive process have been surveyed using genomic DNA polymorphisms in transcribed regions [17,19]. However, the differentiation of the transcriptome between genetic groups has not been addressed yet.

In the present study, we conducted high-throughput transcript sequencing of current-year shoots collected from individuals grown in three common gardens at days when the temperature exceeded 30 degrees Celsius. Three common gardens are located in the north (Miyagi Prefecture), in the central area (Ibaraki Prefecture), and in the south (Kumamoto Prefecture) of Japan. The genetic differentiation between individuals was evaluated using SNPs detected by mapping the RNA-Seq reads to the reference genome [20]. The genetic differences were further compared to the representative gene expression patterns of co-expressed gene modules detected by WGCNA to find the gene modules associated with local adaptation. If the selection pressure from the local environment induces phenotypic divergence, the differences in the expression of certain genes can explain the divergence. In such cases, the adaptive genes expression will show the patterns associated with the genetic distance between individuals or with climatic variation between their origins. This study aims to identify such gene expression divergence associated with local adaptation and the genes that have central role in such module, it will be crucial for the future conservation of *C. japonica*.

## Materials and Methods

### Plant materials

The details related to the collection of individuals from natural populations and the establishment of a nursery are reported previously [19]. Cuttings from the nursery were grown in a field at the Forestry and Forest Products Research Institute (Tsukuba, Ibaraki, Japan) for 2 years. They were then transferred to three common gardens, one in the Ibaraki (IBR) Prefecture, one in the Kumamoto (KMT) Prefecture, and the other in the Miyagi (MYG) Prefecture, which were established based on a randomized design. Approximately 1000 trees were transplanted in each garden in 2015, 2017 and 2014, respectively. Out of the 23 populations grown in these gardens, 12 populations were selected to represent the natural distribution of *C. japonica* (Fig 1, Table 1). Two clones per population (24 clones in total) from each common garden were subjected to transcriptome analysis. Ramets from the same individual were used in all three common gardens when possible.

**Fig 1.**
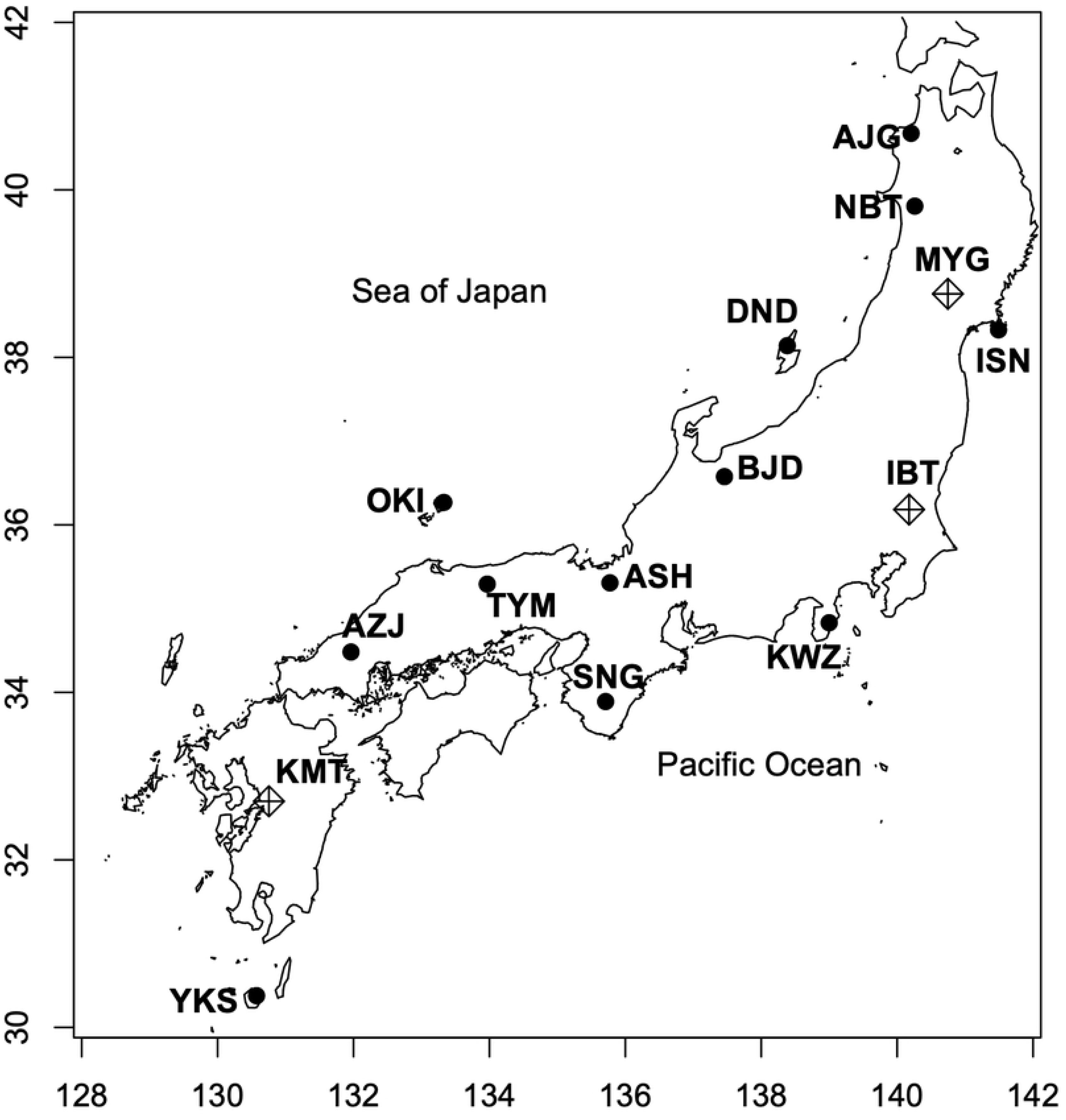
Locations of source populations (denoted by circles) and common gardens (denoted by diamond pluses)

**Table 1.**
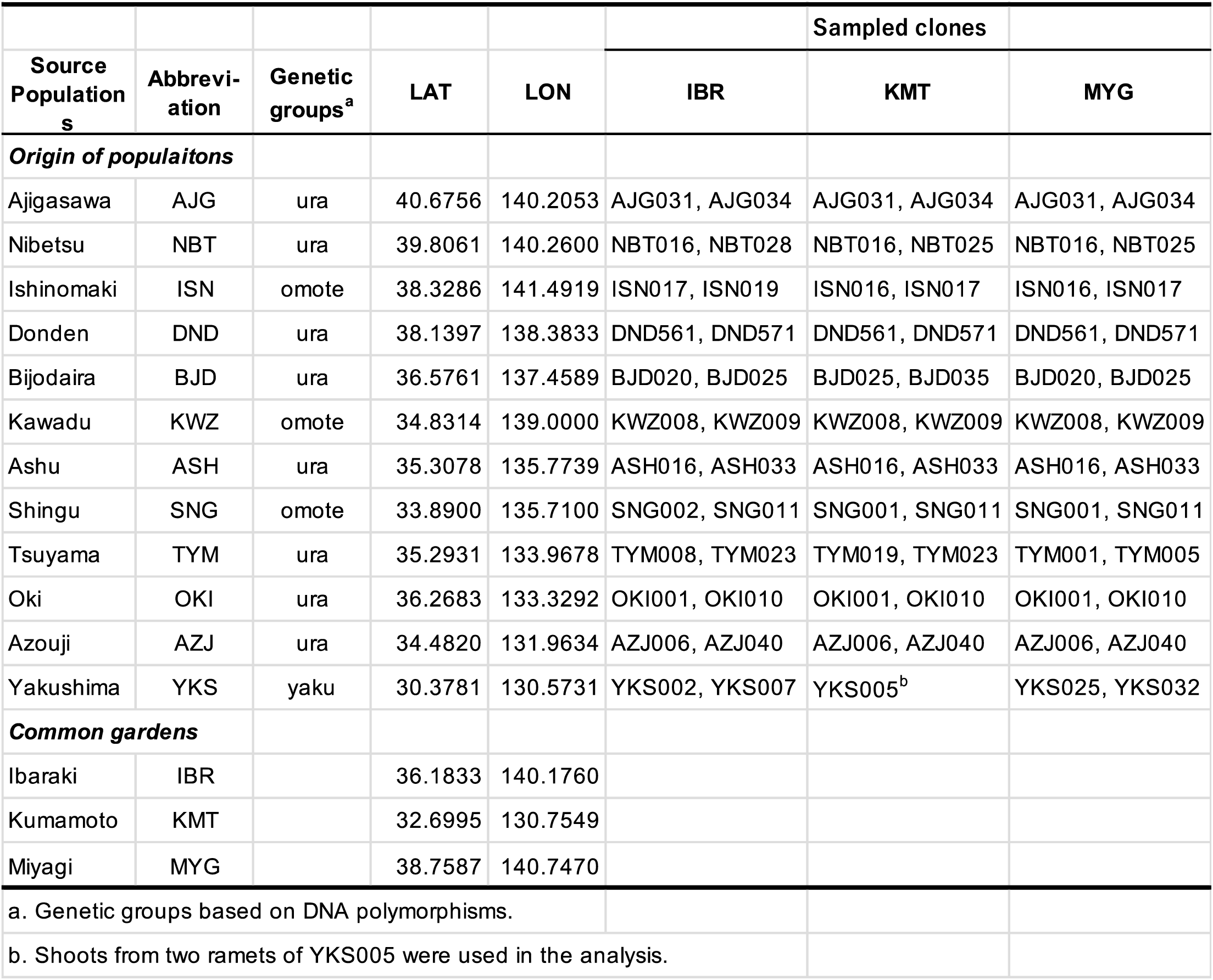
Locations of the analyzed populations and common gardens.

#### Estimation of climate transfer distances

The historical bioclimatic variables for the original locations of populations and the common gardens were extracted from WorldClim at a scale of 30 arc seconds using the R package "raster" [21] (S1 Table). In addition, the bioclimatic variables of the common gardens during the period in which the material trees were grown were calculated using climatic values obtained from Agrometeorological Grid Square Data (AMGSD)[22], using the R package "dismo" [23]. It demonstrated a clear increase in temperature at each site, with an annual average temperature increase of approximately 1°C at each location (TableS1, Fig S1). The differences in climate variables between original locations and common gardens were evaluated via principal component analysis (PCA) using dudi.pca in the R package "ade4" [24]. More climate transfer differences (CTD) would be more stressful conditions to the material trees and thus may shape the gene expression. To assess the hypothesis, CTD between original locations and common gardens were calculated as described in previous study [2], using the top four principal components (PCs) of the PCA.

### RNA-Seq

Current-year shoots of well-sunned branches were sampled on June 28^th^, 2022, at IBR, August 8^th^, 2020, at KMT, and on August 2^nd^, 2021, at MYG. The samples were collected in these hot summer days so that the transcriptome analysis would be able to provide more information on genes involved in the response to high temperatures. Sampling was conducted at around 2pm at all sites to reduce the difference caused by circadian oscillation. The air temperature at this time at IBR, KMT, and MYG was 36℃, 34℃ and 30℃, respectively (S2 Fig). The most recent day of rainfall before sampling was 9 days prior in IBR, 6 days prior in KMT and 2 days prior in MYG. Total RNA was extracted from the shoots using a Maxwell RSC Plant RNA Kit (AS1500) and Maxwell RSC48 instrument (Promega). In addition to being subjected to the protocols required by the kit, the samples were pre-treated with CTAB solution (2% CTAB, 2% PVP, 100 mM Tris-HCI [pH 8.0], 25 mM EDTA, 2 M NaCl, and 2% β-mercaptoethanol). Pair-ended 150-bp reads were obtained using an Illumina NovaSeq 6000 platform (Novogene).

### Constructing a reference transcript set and obtaining expression data

The obtained reads were filtered using Trimgalore v 0.6.7 (https://github.com/FelixKrueger/TrimGalore) to remove low-quality reads. Reads less than 50 nt in length were then removed using the SolexaQA++ v3.1.7.1 LengthSort command [25]. A transcript sequence set was constructed from reads data obtained in this study (reads are under registration process) and reads used in the previous study to improve the quality of assemblage (reads are under registration process). The whole-genome assembly (SUGI ver.1) [20] was used as guidance for constructing transcript assemblage. First, the reads of each sample were mapped to the *C. japonica* reference genome sequence using hisat2 version 2.2.1 [26]. Transcripts were predicted using the resultant bam files by psiclass [27]. The gtf file generated by psiclass was then merged with standard predicted genes (SUGI_1.std.gene.gff3) [20] by stringtie version 2.2.1 [28].The counts for each transcripts were calculated based on the merged gtf file. Sequence of predicted transcripts were extracted by gffread [29]. Newly predicted genes, which were not included in the SUGI_1.std.gene.gff3, were prefixed with "CJHT".

### Annotation of reference transcripts

The completeness of the assembly was assessed by comparing it with the embryophyta_odb10 database including the core gene set of land plants in BUSCO v5 with default parameters [30]. The reference sequences were compared to Araprot11_pep_20220505.fa, which was retrieved from the TAIR database (www.arabidopsis.org) [31], and the UniProtKB/Swiss-Prot database using blastx 2.10.0+ with 1e-5 as the cutoff value [32].

### Single nucleotide polymorphism detection

After filtering any potential PCR duplicates using the rmdup command of samtools v1.13 [33], single nucleotide polymorphisms (SNPs) between samples were extracted via the mpileup command of bcftools v1.16 [33]. The obtained SNPs were further filtered using the filter command of bcftools with options -g 3, -G 10, and -e ‘QUAL≤30||DP<100||MAF≤0.05’ to obtain reliable SNPs. Discordant genotypes between ramet samples from the same individual were excluded using an in-house perl script. SNP frequency per 1kb was calculated for each polymorphic gene. SNPs from the genes with high SNP frequencies (≥10 SNPs per 1kb) were excluded to avoid possible false SNPs caused by paralogous genes or multi-gene families.

### Genetic differentiation among samples

To assess genetic differentiation in the samples examined in this study, PCA was conducted based on SNPs using dudi.pca in the R package "ade4" [24]. One SNP per polymorphic gene was randomly selected for the PCA using an in-house perl script. The results were visualized using the fviz_pca_ind function in the R package "factoextra" [34].

### Overall transcriptome comparison

The correlations of all expression profiles between samples were calculated using the cor function in R with the “spearman” method and visualized using the R package "corrplot" [35]. PCA was also conducted for gene expression data to assess the overall similarity of the transcriptome between samples. To normalize the raw count data, the variance stabilization transformation (vst) was applied via the iDEP website tool [36]. PCA was carried out for the top 1000 transcripts with the highest quartile coefficient of dispersion (QCD) between samples for each common garden using dudi.pca in the R package "ade4" [24].

### Gene expression differentiation detection by WGCNA

To reduce the effect of random fluctuation in low-expression transcripts, transcripts whose expression was lower than median expression value were first removed. We then focused on top 5000 transcripts with the highest QCD among filtered transcripts for each common garden. To construct consensus gene networks of three common gardens, the 5000 transcripts from each site were concatenated and the set consisting of 8286 transcripts was used in WGCNA. The vst counts were used as the expression value of each transcript. WGCNA was conducted using the R package “WGCNA” to survey gene networks that showed expression patterns associated with genetic differentiation [7]. The soft-threshold power for detecting co-expressed networks (gene modules) was set to 6 in constructing a consensus network, and the networks were constructed using the blockwiseConsensusModules function with default settings, except for corType = “bicor,” networkType = “signed,” maxBlockSize = 10000, and maxPOutliers = 0.1. The correlation coefficients between the module eigengene (ME) of each detected gene module and the first, second PCs of the PCA using SNPs (PC1_SNP_ and PC2_SNP_) were calculated using the corr.test function in the R package “psych” with the “spearman” method [37]. The ME is indicative of the characteristics of each module, which gives the most representative gene expression in a module. We selected modules whose MEs showed a correlation with PCs with an adjusted P-value of ≤ 0.1 as modules significantly associated with PCs. The correlation between climatic variables (bio1-bio19, CTD) and the eigengene expression was also calculated in the same way. Candidate hub genes were selected if their module memberships (MM) to the corresponding module were equal to 0.6 or more and the absolute values of their gene significances (GS) were equal to 0.5 or more.

### Gene ontology enrichment analysis

Gene Ontology (GO) enrichment analyses of genes in modules associated with PC1_SNP_ and PC2_SNP_ were conducted using the R package "clusterProfiler."[38]. The reference assemblage used in this study was annotated with Gene Ontology terms based on the annotation of the best matches in *Arabidopsis* reference genes. The annotation list of 8286 transcripts applied in the WGCNA was used as a background list.

## Results

### Constructing reference sequence*s*

The average mapping rates of samples to the reference genome sequence (SUGI v1) were 95.7%, 96.6%, and 96.3% for IBR, KMT and MYG, respectively. The assembly of obtained reads guided by the reference whole-genome sequence resulted in 77,212 transcripts from 56,203 genes. The N_50_ value of the merged reference transcript set was 1,962 bp. The homology search against *Arabidopsis* reference sequences and the UniProtKB/Swiss-Prot database showed that 42,150 genes (75.0%) exhibited homologous sequences in at least one database with a significant threshold (e-value <1e-5). BUSCO analysis showed 93.3% of the 1614 conserved genes in embryophyta_obd10 were present in the constructed transcript set. Specifically, 87.4% were single-copy genes, and 5.9% were duplicate genes.

### SNP detection

A total of 110,869 polymorphic sites were detected in 13,344 genes after filtering out the sites that showed discrepant genotypes between ramets. The 1,983 insertions or deletions found in 1,701 genes were also excluded. The frequency of SNPs in a polymorphic gene ranged from 0.04 to 51.3 per 1 kb. As mentioned in the Materials and Methods section, the SNPs in genes with high SNP frequencies were excluded from further analysis to avoid possible false SNPs due to the misalignment of reads to regions with high sequence similarity, such as paralogs and multi-copy gene families. As a result, 94,732 SNPs in 12,389 genes were retained. The average SNP frequency was one SNP for every 506.4 bp.

The samples were classified into three groups by PCA using SNP genotypes of those 12,389 loci (S3 Fig). The results were consistent with the previously reported classification of genetic groups: ura-sugi, omote-sugi, and individuals originating from Yakushima Island (hereafter referred to as yaku-sugi). Based on the PCA results, we decided to include individuals from the northern peripheral region in ura-sugi in this study. The first axis (PC1_SNP_) separated the three groups and showed yaku-sugi as more genetically differentiated from the other groups, whereas the second axis (PC2_SNP_) separated omote-sugi from the other groups. A total of 40,783 synonymous and 39,911 non-synonymous nucleotide exchanges were found in the coding region of 11,627 and 11,547 genes, respectively. Unique substitutions for specific genetic groups were not detected.

### Overall transcriptome profile of samples from the common gardens

After filtering low-expression transcripts, no transcripts were exclusively expressed in specific populations or genetic groups. There was a positive correlation between the overall transcriptome of the samples, with the correlation coefficients ranging from 0.75 to 0.99 (S4 Fig). Even though they were grown under different environmental conditions, ramets derived from the same individual tended to show more correlated gene expression patterns than a pair of individuals with the same origin grown in the same common garden. On the other hand, the PCA using expression data of highly variable transcripts indicated that expression patterns were associated with the genetic groups in IBR and KMT, whereas such association was not apparent in MYG (Fig 2).

**Fig 2.**
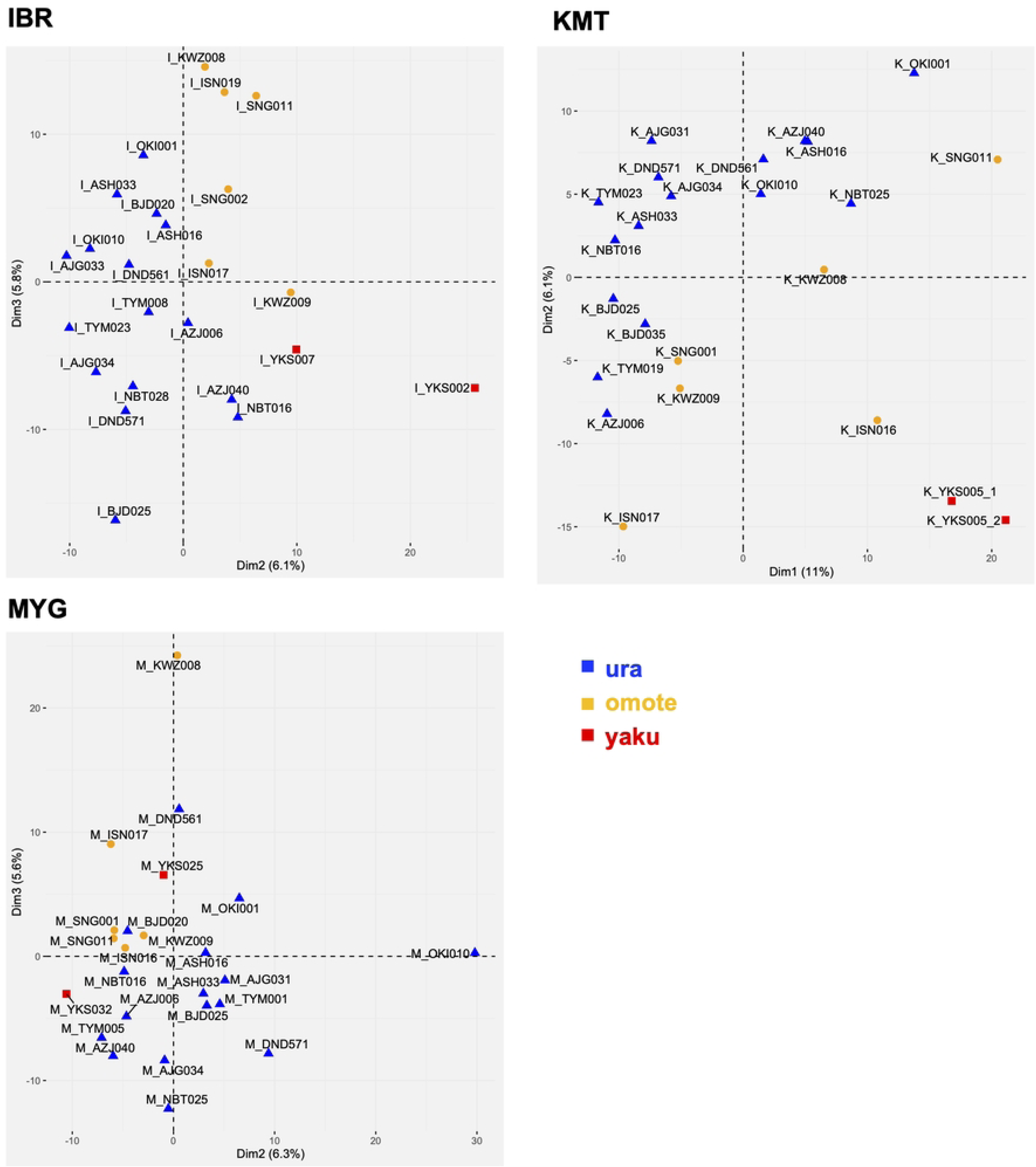
PCA plot of samples based on the expression patterns of 1000 transcripts with high qcd values and specific expression levels (see Materials and Methods)

### Transcripts associated with genetic differentiation

To identify gene modules exhibiting differential expression levels associated with the genetic differentiation, we examined the correlation between SNPs and the MEs of each module. A positive correlation indicated that the gene expression was higher in the omote-sugi, while a negative correlation indicates that the gene expression was higher in the ura-sugi.

A total of 25 co-expressed gene modules were detected by WGCNA, and MEs of three, nine, and two modules were correlated with PC1_SNP_ for IBR, KMT, and MYG site, respectively (Table 2). Among them, ME of grey60 module (MEgrey60) was positively correlated for all three sites, while the MEdarkred was negatively correlated. For PC2_SNP_, three and one modules showed significant correlation in IBR and KMT, respectively, and MEwhite was positively correlated in both sites. No moules were correlated with PC2_SNP_ in MYG.

**Table 2.** Detected gene modules and correlation to PC_SNP_.

| **Table 2** Module eigengene with significant correlation to PC<sub>SNP</sub>. |  |  |  |  |
| --- | --- | --- | --- | --- |
| modules | r<sup>a</sup> | p | FDR | gene numbers |
| **IBR** |  |  |  |  |
| PC1<sub>SNP</sub> |  |  |  |  |
| MEgrey60 | 0.63 | 0.0010 | 0.0400 | 59 |
| MEdarkred | -0.64 | 0.0008 | 0.0400 | 43 |
| MEgreen | -0.55 | 0.0060 | 0.0900 | 395 |
| PC2<sub>SNP</sub> |  |  |  |  |
| MEwhite | 0.56 | 0.0046 | 0.0900 | 36 |
| MEgreenyellow | 0.56 | 0.0047 | 0.0900 | 152 |
| MEpurple | 0.54 | 0.0070 | 0.0900 | 153 |
| **KMT** |  |  |  |  |
| PC1<sub>SNP</sub> |  |  |  |  |
| MEgrey60 | 0.78 | 0.0000 | 0.0005 | 59 |
| MEturquoise | 0.68 | 0.0002 | 0.0080 | 1646 |
| MEpurple | 0.58 | 0.0031 | 0.0400 | 153 |
| MEmagenta | 0.54 | 0.0061 | 0.0680 | 164 |
| MEtan | 0.53 | 0.0072 | 0.0700 | 133 |
| MEorange | 0.51 | 0.0086 | 0.0820 | 37 |
| MEred | -0.66 | 0.0004 | 0.0100 | 615 |
| MEdarkred | -0.64 | 0.0008 | 0.0150 | 43 |
| MEsalmon | -0.62 | 0.0012 | 0.0200 | 87 |
| PC2<sub>SNP</sub> |  |  |  |  |
| MEwhite | 0.51 | 0.0104 | 0.0810 | 36 |
| **MYG** |  |  |  |  |
| PC1<sub>SNP</sub> |  |  |  |  |
| **MEgrey60** | 0.69 | 0.0002 | 0.0163 | 59 |
| **MEdarkred** | -0.66 | 0.0004 | 0.0163 | 43 |
| CTD |  |  |  |  |
| MEgrey60 | 0.58 | 0.0819 | 0.0819 | 59 |
| a. Only those with the absolute value of a correlation coefficient of 0.5 or more are shown. |  |  |  |  |

### Gene enriched in modules exhibiting common correlations to genetic differences at several test sites

For grey60 module, which was positively correlated PC1_SNP_ in three common gardens, no Gene Ontology (GO) term was enriched (S2 Table). However, manual inspection of gene annotation in this module showed 24 genes related to disease resistance (S3 Table). Two genes (CJHT.4193 and CJHT.3914) were selected as hub-genes in all common gardens; one showed similarity to the retrotransposon-like region of Arabidopsis CRK8 (AT4G23160) and another has sequence similarity to AT2G38430 (S4 Table).

On the other hand, darkred module that was negatively correlated with PC1_SNP_ was enriched with terpenoid biosynthetic genes (S2 Table). Despite the absence of genes that met the requisite criteria of hub-genes across all three common gardens, genes with sequence similarity to Arabidopsis terpene synthase 03 (AT4G16740) demonstrated a robust correlation with the other genes and with PC1_SNP_ (S4 Table).

White module, which was correlated with PC2_SNP_ in both IBR and KMT, was enriched with "positive regulation of flavonoid biosynthetic process" (S2 Table). Eight of the genes met the criteria for hub genes, with one being homologous to Arabidopsis RPM1 (AT3G07040), which is involved in disease resistance (S4 Table). The remaining seven genes exhibited no homology to any known genes. Notably, the sequences of these seven genes displayed a high degree of sequence identity (S5 Fig), and furthermore, all of the hub-genes, including RPM1, were located within an adjacent region of chromosome 7 of the reference genome sequence (S4 Table)[20].

### Heat and light response genes excessed in omote-sugi at IBR

In addition to grey60 and darkred modules, green module correlated with PC1_SNP_ in IBR, which enriched "signal transduction". For PC2 _SNP_, two more modules (greenyellow and purple) in addition to white were positively correlated in IBR. MEs of these modules have a similar expression pattern (S6 Fig), and both are enriched with genes responding to heat and oxidative stress (S2 Table). In addition to those GO terms, greenyellow module was characterised by a high number of genes related to light response (S2 Table).

### More modules correlated to genetic differentiation at KMT

In KMT, an additional five modules to grey60 have positive correlation with PC1_SNP_ (Table 2). Enriched gene ontology term indicated that turquoise module containing many photosynthesis-related genes, magenta module containing defense response genes, tan module containing lipid metabolism genes, whereas no GO term was enriched in orange module (Table2 and S2 Table). On the other hand, MEsalmon and MEred were negatively correlated with PC1_SNP_. Salmon module was enriched with lipid biosynthetic related genes. Although no GO term enriched in red module, both modules’ hub-genes included GDSL-like Lipase and gibberellin biosynthesis genes (S4 Table). The gene expression patterns of the two modules, however, did not exhibit high similarity at KMT (S6 Fig).

### Transcripts associated with climate variables

The correlation between the climatic variables (bio1-19) and the eigengene expression of modules were also assessed to explore what environmental conditions might have affected gene expression. More temperature-related variables show a high correlation with the eigengene expression than precipitation-related variables (S5 Table), especially temperature seasonality (bio4) and the annual range of air temperature (bio7). Only MEdarkred correlated with seasonality of precipitation. MEturquoise showed a highly significant correlation to the mean daily minimum air temperature of the coldest month (bio6) at KMT (S7 Fig). Correlation with CTD was detected only with MEgrey60 in MYG.

## Discussion

This study employed a comparative transcriptomic approach to examine the genetic differentiation of *C. japonica* individuals grown in three common gardens exhibiting different environments in the north, the center, and the south of Japan. The transcriptomes provide information on both genetic polymorphism and gene expression differences that exist between natural populations of different geographical origins. It is challenging to ascertain whether the genetic polymorphisms are the consequence of functional selection or whether they emerge by chance due to geographic distance. In contrast, gene expression differences correlated with genetic or climatic differences have more likely to be associated with regional adaptation of the species, because changes in gene expression have more likely to result in phenotypic differentiation. Overall, gene expression profiles were more similar between individuals grown at different common gardens than individuals in the same genetic groups (i.e. omote, ura, yaku-sugi) grown at the same common garden (S4 Fig). The results indicate that gene expression in each individual is strongly influenced by genetic factors, despite substantial variation observed within the same genetic groups, even among individuals from the same origin (Fig 2). This may also suggests that the expression of most genes analyzed has not been altered by epigenetic regulation, despite the different environmental conditions between the common gardens (S1 Fig).

The expression of the genes with higher variability between samples, however, was more similar within genetic groups at IBR and KMT (Fig 2), whereas no such tendency was observed at MYG. The air temperature on the sampling date at IBR and KMT was higher than at MYG (S2 Fig). In addition to the temperature on the sampling date, the IBR and KMT test sites were likely drier than the MYG (S2 Fig). One possibility is that the expression of genes responding to higher temperatures or drought might be differentiated between genetic groups.

### Omote-sugi had higher expression of disease resistance genes

In the present study, we were able to identify gene modules whose expression levels correlated with indicators of genetic differentiation as determined by single-nucleotide polymorphisms (SNPs). A positive correlation was observed between PC1_SNP_ and the expression of defense response genes, while a negative correlation was detected between PC1_SNP_ and the expression of terpenoid biosynthesis genes in three common gardens. These correlations detected at the three common gardens in different part of Japan suggests that there may be fixed genetic changes between genetic groups that led to the changes of gene expression.

The grey60 module, which include many disease-resistance genes, highly expressed in omote-sugi. The hub gene of this module is similar to the retrotransposon-like region of *CRK8* of Arabidopsis. Although the exact function of this gene has not been determined, an association between polymorphisms in the retrotransposon-like region of *CRK8* and quantitative disease resistance to the fungal pathogen *Sclerotinia sclerotiorum* was previously reported [39]. The author speculated that the retrotransposon region is not actually transcribed, but a corresponding region is likely transcribed in *C. japonica*. The region includes a Reverse transcriptase- and Ribonuclease H-like domains, and is conserved among a wide range of plant species. The sequences with similarities to *CRK8* are distributed in the *C. japonica* genome at a relatively high frequency. The role of this gene in disease resistance is highly intriguing. The function of another hub-gene detected in this module, AT2G38430, also remains unclear. However, a highly similar gene, AT3G54310, is induced by bacterial quorum sensing (QS) molecules [40]. It is suggested that this hub-gene is also involved in disease resistance.

In addition to grey60 module, white module is also related to a disease response, since the hub-gene is homologous to *Arabidopsis* RPM1. RPM1 is the NB-ARC domain-containing disease resistance protein that is involved in effector-triggered immunity [41]. The module was positively correlated with PC2_SNP_ that separates omote-sugi from other genetic groups, at the IBR and KMT sites. In IBR and KMT, expression pattern of MEwhite was similar to MEgrey60 (S6 Fig). Thus, the functional interaction may exist between these two modules. As described earlier, the hub-genes in white module were located in an adjacent region of chromosome 7. The tight linkage of transcription patterns and the physical position within the genome suggests the importance of these novel genes in defense response. Furthermore, magenta module enriched with defense response genes, especially systemic acquired resistance related genes, moderately correlated to the PC1_SNP_ at KMT site.

The excess of modules enriched with defense response genes in omote-sugi can be due to gene induction caused by pathogens that preferentially attack them. Correlation of MEmagenta to PC1_SNP_ observed only at KMT might reflect such specificity. On the other hand, genes in grey60 module were expressed more in omote-sugi than ura-sugi at all three common gardens. As previous study demonstrate [14], regional variations of pathogens associated with *C. japonica* are present throughout Japan. It is therefore that the pathogens present at three experimental sites are more likely to be diverged. Warm winters and spring and summer precipitation have been shown to be associated with the abundance of Swiss needle cast (SNC) in Douglas fir [42]. Summer humidity and annual temperatures tend to be high in the areas where omote-sugi is distributed. This might make the pathogen more active in those regions, and an adaptation may have occurred in omote-sugi to constitutively express more genes against biotic stress at summer.

### Ura-sugi had higher expression of terpenoid biosynthesis genes

The darkred module involved terpenoid biosynthesis genes exhibited a negative correlation with PC1_SNP_ and thus were highly expressed in ura-sugi. Terpenoids have a wide range of roles in responding to biotic and abiotic stress [43]. Although terpenoids have function in protection from high temperature [44], genes included in the module more related to biotic stress (S4 Table). The stored and volatile terpenoids of *C. japonica* were examined in the summer of 2019 for 12 populations at MYG, including all those analyzed in the present study except for BJD and TYM [14]. In that study, the amount of stored terpenoids was negatively correlated with warm and less snowy climates. It is consistent with the results obtained in this study, but the volatile terpenoids showed different geographic patterns. The candidate hub gene for this module exhibits sequence similarity to Arabidopsis *TSP03*, which has been demonstrated to play a role in the variation of herbivore-induced volatile terpenes among ecotypes. The amount of volatile terpenes may be related to the stored terpenes, but the functional meaning of observed difference in the gene expression needs further investigation. Mapping of RNA-Seq reads predicts the existence of multiple transcripts of this locus. In *Arabidopsis*, sequences of *TPS03* transcripts and their functions also diverged between ecotypes [45]. The multiple transcripts detected in this study may also reflect functional divergence among genetic groups.

As mentioned earlier, regional differences in biotic stress may have adjusted the gene expression. The observed results may also be explained, however, by the growth-differentiation balance hypothesis [46]. The hypothesis states that plants with slower growth rates tend to produce more secondary metabolites. Uras-ugi is known to grow more slowly than omote-sugi at initial growth [16]. Although the available data are limited to the early growth stages, the growth measurements conducted at KMT and MYG common gardens also indicate a trend toward slower growth in ura-sugi. Ura-sugi might have evolved to produce more secondary metabolites, such as terpenoids, in order to exhibit constitutive tolerance to biological stresses.

### Differentially expressed abiotic stress response genes between genetic cluster

As noted above, there was evidence of constitutive differentiation in the gene expression associated with biotic stresses between genetic groups, but differences in gene expression in response to abiotic stresses were also observed at specific common gardens.

At IBR, genes responding to heat and oxidative stress were highly expressed in omote-sugi (MEgreenyellow, MEpurple). The sampling at IBR was done in June, the beginning of the summer when temperatures were first on the rise. It is possible that the individuals had not been acclimated to high temperatures and perhaps high light intensity at the sampling date. This may have resulted in the induction of genes that respond to heat and light stress at this site (S2 Table). The hub-genes of greenyellow module were mainly small heat shock proteins, *HSP17.6C (AT1G53540)* and *HSP21 (AT4G27670)*, both are known to be involved in heat acclimation and to be epigenetically regulated [47,48]. The observed difference, however, cannot be attributed to differences in temperature of origin because no association was detected between the temperature-related climate variables and the eigengene expression of these modules. The selective pressure to make higher expression in omote-sugi was thus unclear. Yaku-sugi was previously assumed to the omote-sugi lineage, but the amplicon sequencing of 120 nuclear genes proposed the classification of yaku-sugi as a distinct variant in a lineage that prior to the divergence of both ura- and omote-sugi lineages [49]. The observed gene expression difference yaku- and omote-sugi gene modules associated with PC2_SNP_ further support the hypothesis of divergence between these two groups. At the KMT site, turquoise module, enriched with genes respond to reactive oxygen species and photosynthesis, tend to express higher in yaku- and omote-sugi (Table 2, S7 Fig). The plants adjusted their photosynthetic characteristics to their growth temperatures [50]. There were high correlations between the MEturquoise and the bio-climatic variables related to the temperature (S5 Table, r>=0.7 for bio4, bio6, bio7, and bio11, and r>=0.6 for bio1). The photosynthetic process genes of individuals from warmer regions may be adapted to high temperatures, whereas individuals from cooler regions may lack such adaptations. These differences in temperature adaptation are likely to contribute to the observed variation in gene expression at KMT. Contrary to our expectation, however, there was no correlation with bio5, which measures temperature in the warmest season (summer). Given the relatively limited variation in bio5 values across different regions (S1Table), it is plausible that the regional adaptation of photosynthesis might be influenced by winter temperatures.

Addition to the modules showed higher expression in omote-sugi, the two modules (salmon and red) were expressed higher in ura-sugi at KMT. As mentioned earlier, these two modules include many GDSL-like Lipase/Acylhydrolase superfamily proteins and gibberellin biosynthesis-related proteins. Although their MEs were not correlated at KMT, they showed similar expression patterns at IBR and MYG (S5 Fig). GDSL-like Lipases are a large family in plant species, and are related to developmental regulation and response to both biotic and abiotic stresses [51]. In other plants, the regulation of GDSL-like Lipases by gibberellin suggested by the sequencing analysis of the promoter regions [52]. Further dissection of the function of these genes will contribute better understanding for the role of gibberellin and GDSL-like Lipases in environmental adaptation.

## Conclusion

A previous study [19,20] identified several environmentally adaptive genes involved in the differentiation of omote-sugi and ura-sugi, but they did not match the hub-genes in modules associated with genetic differentiation detected in the present study. It should be noted that the results presented here were based on limited data. Transcriptome analysis was conducted only once at each common garden, and the number of individuals analyzed was not particularly large. To understand the environmental adaptation of *C. japonica* trees, it is necessary to obtain data from many individuals at more time points. Nevertheless, this study demonstrated the usefulness of transcriptome analysis in the field to find the candidates of adaptive genes. The results suggested that the two main genetic groups have a differentiated regulation of gene expression in response to mainly biotic stresses. Due to climate change, these stresses will increase at a more rapid pace compared to the ability of *C. japonica* to adapt to them, potentially disrupting the existing adaptive relationships [53]. The adaptation of *C. japonica* to biotic stresses, in addition to abiotic stresses, is a critical factor to consider future conservation efforts of this species.

## Acknowledgments

The author thanks Ms. Furusawa for her excellent assistance with the experiments. The authors are grateful to Y. Matsui, K. Yokoo, and R. Kusano from Kumamoto Prefecture Forestry Research and instruction Center, T. Sasaki and K. Sato from Kawatabi Field Center of Tohoku University for the maintenance and preparation of research materials.

## Data Archiving Statement

Registration to the DDBJ DRA database is in progress for the 72 Illumina sequence read data reported in this manuscript, and the accession numbers will be supplied once available.

## Figure legends of Supplemental materials

**S1 Fig.** Climatic differences between the locations of source populations and common gardens. PCA is based on 19 climatic variables retrieved from WorldClim or calculated based on the current climate data retrieved from AMGSD (denoted by _C, Ohno et al., 2016)

**S2 Fig.** Weather conditions for 30 days before sampling. (a) max air temperature of the day, (b) total precipitation of the day. Weather data for each common garden obtained from Agrometeorological Grid Square Data (AMGSD; Ohno et al., 2016)

**S3 Fig.** Principal components analysis using SNP of 12,389 loci between analyzed 35 samples. One SNP per locus was randomly selected. Three genetic groups were shown in blue triangle (ura-sugi), yellow circle (omote-sugi), and red square (yaku-sugi).

**S4 Fig.** Correlation of overall expression profile between samples based on variance stabilized formation counts. Darker green indicates a higher correlation.

**S5 Fig.** Sequence identities between candidate hub genes of MEwhite except for SUGI_0703660.

**S6 Fig.** Hierarchical clustering dendrogram of module based on their expression patterns.

**S7 Fig.** Correlation between the eigengene expression of MEturquoise in KMT and bio6 (mean daily minimum air temperature of the coldest month).

